# Glioblastoma-derived extracellular vesicles released after radiation promote cognitive impairment through NFκB-mediated microglial activation

**DOI:** 10.64898/2026.06.09.730969

**Authors:** Sara Macias Palacio, Nicole Rummel, James Campbell, D. Allan Butterfield, Subbarao Bondada, Chi Wang, Abu Saleh Mosa Faisal, John Villano, Bjoern Bauer, Daret St. Clair, Luksana Chaiswing

## Abstract

Glioblastoma (GBM) is the most aggressive primary brain tumor in adults. Cognitive impairment is a common sequela in glioblastoma survivors, yet the underlying mechanisms remain poorly understood. Extracellular vesicles (EVs) derived from glioblastoma are established mediators of intercellular signaling within the tumor microenvironment. Here, we investigated whether GBM-derived EVs released after radiation treatment (RT-EVs) regulate cognitive function. Treatment with RT-EVs was associated with cognitive deficits and neuroinflammatory responses in vivo. In vitro, RT-EVs activated the NFκB pathway and induced the release of neurotoxic H_2_O_2_. Importantly, NFκB p50 knockdown abolished the H_2_O_2_ release previously triggered by RT-EVs, demonstrating mechanistic dependence on NFκB signaling. Collectively, these findings identify GBM-derived RT-EVs as critical mediators of cognitive impairment through NFκB-dependent redox imbalance. EV-driven redox dysregulation may therefore represent a therapeutic target to mitigate GBM-associated cognitive dysfunction.

**Highlights:**

- Radiation induces the release of glioblastoma-derived EVs that are biologically different from those released under non-irradiated conditions.
- EVs released from glioblastoma after radiation are sufficient to impair cognition
- EVs from irradiated glioblastoma can activate microglia via NFκB and induce production of neurotoxic H_2_O_2_

**Graphical Abstract:** 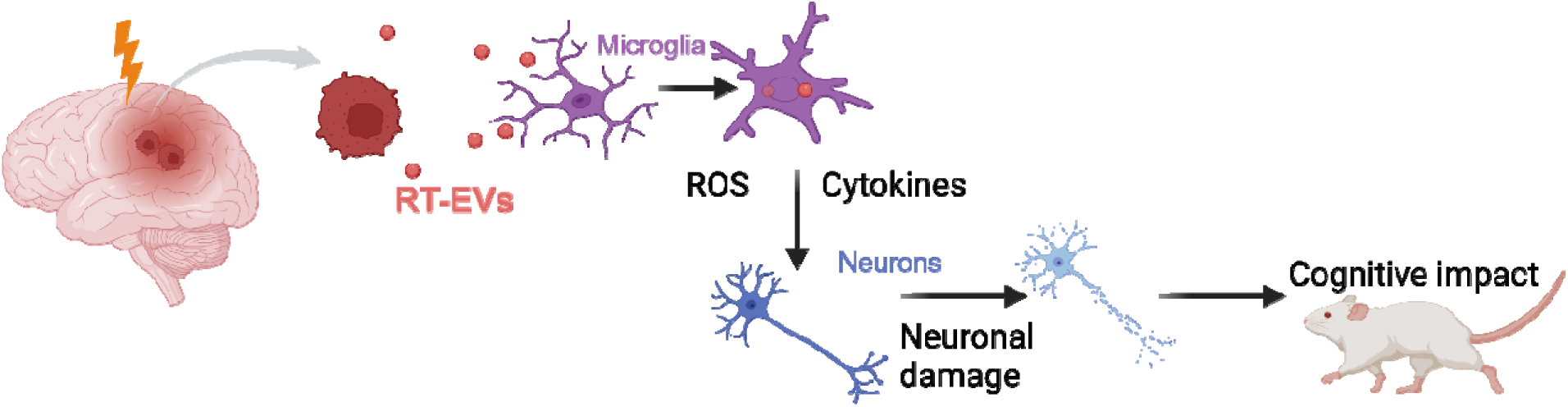

## 1. INTRODUCTION

Over the past three decades, the global incidence and mortality of brain cancer have increased, affecting both sexes and most age groups (Ilic and Ilic 2023). Approximately 80% of these cases correspond to adult-type diffuse gliomas, including astrocytoma (IDH-mutant), oligodendroglioma (IDH-mutant, 1p/19q-codeleted), and glioblastoma (IDH-wildtype), as defined by the current WHO classification of central nervous system tumors (Torp, Solheim et al. 2022; Lennartz, Tholke et al. 2023).

Glioblastoma (GBM) is the most common and aggressive primary brain tumor in adults. Because of its marked aggressiveness, intratumoral heterogeneity, and invasiveness, the current standard of care remains insufficient for durable disease control, despite maximal safe resection followed by radiation and chemotherapy (Lennartz, Tholke et al. 2023; Houskamp, Skorich et al. 2025). This is reflected in the poor prognosis of patients with GBM, whose median survival remains approximately 16 months even after optimization of available therapies (van Solinge, Nieland et al. 2022).

In addition to the poor survival associated with GBM, patients tend to have neurological symptoms like headaches, vision or balance alterations and cognitive decline, including speech difficulties and memory loss, highlighting the significantly lower quality of life of GBM patients compared to those with other types of cancer (Iyer, Ariwodo et al. 2025, Xie and Wang 2025). These deficits can be present before therapy and may worsen during treatment, substantially compromising quality of life and functional independence. These therapy-related cognitive effects are not fully understood, the outcomes in neuropsychological assessments indicate mixed trends, with some cognitive domains showing improvement prior to the treatment while others display better results afterwards (Habets, Kloet et al. 2014, van Loenen, Rijnen et al. 2018).

Understanding treatment responses and secondary effects is complex, considering they are influenced by multiple factors. A key reason is oxidative stress, which is an imbalance of antioxidants and oxidants like reactive oxygen species (ROS) that over time, can cause permanent damage to biomolecules (Albano, Gagliardo et al. 2022). 4-hydroxynonenal is a product of lipid peroxidation that has been proposed as a biomarker for oxidative stress. It is highly reactive and adducts to proteins, altering their function, homeostasis and regulating key molecular pathways (Butterfield and Halliwell 2019, Liu, Cui et al. 2020, Milkovic, Zarkovic et al. 2023, Perluigi, Di Domenico et al. 2024). Nuclear Factor-κB (NFκB) is a family of transcription factors that regulates immune responses (including cytokine release), proliferation, apoptosis, among others. Importantly, the NFκB pathway has been shown to be redox sensitive shaping immune response around the redox status of the cell (Pantano, Reynaert et al. 2006).

Extracellular vesicles (EVs) are membrane-enclosed nanometric particles released by most cells to the extracellular space (van Niel, D’Angelo et al. 2018). As carriers of proteins, lipids, metabolites, and nucleic acids, EVs can reshape recipient-cell behavior locally and at a distance, and they are increasingly recognized as major mediators of intercellular communication in cancer. Growing evidence highlights the role of EVs as biomarkers, modifiers of the tumor microenvironment and mediators in intercellular communication (Urabe, Kosaka et al. 2020, Elias, Hadjiyiannis et al. 2025). However, their impact on cognition and brain function in the context of glioblastoma is yet to be determined.

Given that GBM patients exhibit higher number of EVs in circulation when compared to healthy counterparts as supported by our results and previous reports (Ricklefs, Wollmann et al. 2024), and considering radiation therapy is still given as part of the standard of care for GBM (Obrador, Moreno-Murciano et al. 2024), we aim to explore the role of GBM-derived EVs after radiation in mediating therapy-related cognitive decline.

In this study, we first assessed the effects of radiation on GBM-derived EVs. Then, we found that GBM-derived EVs after radiation cause behavioral and cognitive alterations in vivo. Mechanistically, these EVs can activate microglial cells via NFκB, which in turn secrete hydrogen peroxide that causes neuronal death. Overall, our findings indicate that GBM-derived EVs and their redox and neuroinflammatory effects could be therapeutic targets to improve patient cognition and overall quality of life.

## 2. METHODS

### 2.1. Human Sera Samples

Human serum samples were obtained from adult patients under exempt protocols approved by the appropriate Institutional Review Board (IRB) and in accordance with institutional and federal ethical guidelines. Serum from healthy donors was included as a control group where indicated. Written informed consent was obtained from all participants prior to sample collection. Peripheral blood samples (1 ml/tube) were collected at designated clinical time points as part of standard-of-care procedures. Serum was isolated according to established clinical laboratory protocols and aliquoted into individual vials (one vial per requested time point). Samples were de-identified, assigned unique study codes, and stored under liquid nitrogen until analysis. Associated clinical and demographic data were collected and maintained in a secure database. All specimens were rendered anonymous prior to experimental use to ensure patient confidentiality.

### 2.1. Animals

Animal procedures were approved by the Institutional Animal Care and Use Committee of the University of Kentucky (2021-3793). Adult female C57BL/6 Tyr- mice (10-14 weeks) were purchased from The Jackson Laboratory. All animals were acclimated for a week before any experiments and housed under controlled temperature, humidity and a 14/10 light/dark cycle.

### 2.1. Tissue and blood collection

Blood was collected via cardiac puncture under deep anesthesia. The heart was accessed and blood was aspirated from the left ventricle using a needle. Subsequently, brains were isolated, rinsed in cold PBS and processed for further analyses.

### 2.2. EV isolation

Patient-derived EVs were isolated from sera using the ExoQuick kit (System Biosciences, Palo Alto, CA, USA). Briefly, serum was centrifuged, combined with the ExoQuick solution and incubated for 30 min at 4 °C. The mixture was then centrifuged at 1,500 g for 30 min, supernatant was removed and the pellet was recentrifuged at 1,500 g for 5 min. Finally, supernatant was aspirated and the EV pellet was resuspended.

Cell-derived EVs were isolated from culture supernatant. Cell culture media was filtered through a 0.8□ μm filter. Filtered supernatant was used to isolate EVs utilizing the ExoEasy kit (Qiagen, Germantown, MD, USA) following manufacturer’s instructions. Briefly, filtered media is mixed with binding buffer in a 1:1 ratio. The mix is centrifuged at 500 g for 1 min at RT. Then, wash buffer is added and the mix is spun at maximum speed (up to 5000 g) for 5 min at RT. Finally, EVs are eluted by adding elution buffer, waiting 1 min and centrifuging at 500 g for 1 min. Eluate is placed back on the filter and centrifugation is repeated at maximum speed for 5 min at RT. Isolated EVs are then concentrated using the Amicon Ultra 0.5ml centrifugal filters Ultracel 3k. 450 ul of EV solution is placed on the filter and centrifuged at 14,000 g for 50 min at 4 °C. This is repeated until all solution is filtered. Finally, the filter is flipped upside down to a new elution tube and centrifuged at 14,000 g for 5 min at 4 °C.

### 2.3. Nanoparticle Tracking Analysis

Particle concentration and size of EVs were measured with a ZetaView Nanotracking Analyzer (Particle Metrics, Munich, Germany). The instrument was calibrated with standard beads (110 nm). The EV preparation was diluted in ultrapure PBS at a ratio of 1:10,000. One ml of diluted EVs were loaded in the instrument and rinsing with PBS was performed between measurements.

### 2.4. Immunogold labeling

EVs were fixed in 2% paraformaldehyde in 0.1M sodium phosphate buffer overnight at 4 °C. 5 µl of fixed EVs were placed on a 300-mesh carbon/formvar coated nickel grid, covered for 30 min. Then the grid was washed, transferred to blocking buffer and then incubated with the 4HNE specific antibody (ab48506, Abcam, Boston, MA, USA) overnight at 4 °C. After that, the grid was washed and incubated with the secondary antibody conjugated with 5 nm gold particles (AURION, Wageningen, The Netherlands) for 1 h at room temperature. Finally, the grid was washed, transferred to stabilization solution, dried and stained with 50 ul 1% uranyl acetate in water for 10 min. One final drying step was performed before imaging.

### 2.5. Transmission Electron Microscopy

Imaging was performed as described in (Miller, Xu et al. 2022). Briefly, EVs were fixed overnight with glutaraldehyde at 4 °C. Then, fixed EVs were placed on a copper grid, stained with 2% uranyl acetate, dried and observed under the transmission FEI Talos F200X TEM at 80 kV.

### 2.6. Protein expression via Jess, automated Western Blotting

Cell and EV lysates were extracted using RIPA buffer at 4 °C for 30 min and then centrifuging at 14,000 g for 10 min at 4 °C. Protein concentration was measured using BCA Rapid Gold (Thermo Fisher Scientific). Protein expression was assessed using the Jess instrument (ProteinSimple by Bio-Techne, Minneapolis, MN, USA), a capillary-based automated western blotting system. This was used according to manufacturer instructions using the Simple Western 12-230 kDa separation module with total protein normalization. To visualize and quantify the signal for each protein the Compass software (ProteinSimple by Bio-Techne) was used. Antibodies were diluted at 1:20. The following antibodies were used: 4HNE (ab46545, Abcam, Cambridge, UK), CD68 (ab125212, Abcam), CD9 (sc-59140, Santa Cruz Biotechnology, Dallas, TX, USA), Flippase (HPA052935, Sigma Aldrich, St. Louis, MO, USA), NeuN (NBP1-92716, Novus Biologicals, Centennial, CO, USA), p50 (sc-8414, sc-7178, Santa Cruz Biotechnology), Neurofilament (ab223343, Abcam).

### 2.7. Proteomics analysis using LC-MS/MS

Proteins from EVs were precipitated using trichloroacetic acid and deoxycholate. Then, proteins were enzymatically digested using the EasyPep MS Sample Prep kit (Thermo Fisher Scientific, Waltham, MA, USA) following manufacturer’s instructions. Resulting peptides were separated using liquid chromatography-tandem mass spectrometry (LC-MS/MS). Analysis was performed on a Thermo Scientific Orbitrap Exploris 240 mass spectrometer with an EASY-Spray source housing and a Vanquish Neo UHPLC system (Thermo Scientific). Data were collected using a data-dependent acquisition strategy with positive ionization including stated 2-4 and top N precursor ions in a 3 s cycle. Data were analyzed against the Homo sapiens UniProt database (downloaded March 2024) using the Thermo Scientific Proteome Discoverer (version 3.1) the SEQUEST search algorithm. Downstream data analysis was conducted using proteomics pipelines.

### 2.8. Intranasal administration

Animals were lightly anesthetized and placed with their heads at a 30-45 ° angle. A micropipette was used to deliver a total of 1.5 x 10^10^ per mouse. EVs diluted in 20µl of buffer (10 µl per nostril). A brief interval was allowed between aliquots to prevent overflow.

### 2.9. Microglia and neuron isolation from the adult mouse brain

Brains were isolated as previously described and dissociated using the adult brain dissociation kit and the gentleMACS Octo Dissociator following manufacturer’s recommendations. Then, the solution was filtered with a MACS SmartStrainer (70 µm) and debris and red blood cell removal was performed. CD11b microbeads were added to target microglia. The solution was then separated using MACS column under a magnetic field. The positive fraction was washed and collected as the microglial population. The negative fraction was treated with non-neuronal cells biotin-antibody cocktail, incubated, washed and resuspended. After that anti-biotin microbeads were added, incubated and magnetic separation was performed with the MACS column. Unlabeled cells (negative fraction) were collected as the neuronal population. All the reagents for brain dissociation, microglial and neuronal isolation were purchased from Miltenyi Biotec (Bergisch Gladbach, Germany).

### 2.10. Intracranial administration

EVs were prepared, kept sterile and on ice. Animals were injected with buprenorphine (1 µl/g) and placed on isoflurane chamber. Once anesthesia reached desired depth, animals were secured on a stereotactic frame (Stoelting, Wood Dale, IL, USA) and anesthesia switched to nose cone. Eye ointment (Alcon Laboratories, Fort Worth, TX,USA) was applied. The previously shaved scalp was disinfected alternating with 2% chlorhexidine and saline. A 1 cm sagittal incision was made starting at the eyes, then the skull was cleaned with 3% hydrogen peroxide and 70% ethanol. A micromotor high-speed handheld drill (Stoelting) was used to drill through the skull (3 mm right of the sagittal suture and 3 mm anterior to the lambda). EVs were resuspended and 2.5 x 10^9^ EVs were loaded into a Hamilton syringe (5 µl, 26ga). The syringe was inserted in the hole and EVs were injected at a rate of 1µl/min using a quintessential stereotaxic injector (Stoelting). The syringe was left in the hole for a minute before pulling it out, closing the hole with bone wax (Braintree Scientific, Braintree, MA,USA), cleaning the incision with saline and stapling the scalp.

To mimic EV release from a tumor, the EV administration was performed on the right side because female GBM patients have higher incidence on the right lobe of the brain (Carrano, Juarez et al. 2021).

### 2.11. Behavior tests

Mice were habituated in the test room for 30 min before every experiment. All arenas were disinfected before each trial and between animals. Behavior was recorded using an overhead camera and then analyzed using EthoVision.

#### Novel Object Recognition (NOR)

Each arena was 50 x 50 cm, with an opaque floor and illuminated at 25 +/-3 lux and a background noise of 45 db. Every arena had visual cues in two of the walls for spatial orientation. During the AA phase mice were placed in the arena for 10 min with two identical objects. During the AB phase mice returned to the arena for 10 min but one of the objects was replaced for a novel object. Exploration was defined as the mouse being less than 2 cm from the object. Recognition memory was quantified using the discrimination index, calculated as:

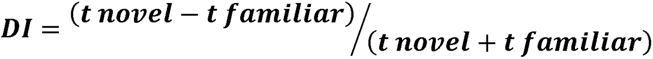

where *t novel* is the time spent exploring the novel object and *t familiar* is the time exploring the familiar object.

#### Fear Conditioning (FC)

Each conditioning chamber was equipped with a grid floor for shock delivery. On day one mice were placed in the chamber with context A (striped walls, disinfectant smell, grid floor) and allowed to explore for 3 min. Then, 5 pairings of an auditory tone (80 dB, 14 s) and a foot shock (0.5 mA, 1 s). 24 h after mice were re-exposed to context A during 3 min without tone or shock. Then, mice were placed in context B (white walls, vanilla scent, black floor) and allowed to explore for 3 min followed by three tones. For all stages the freezing behavior was quantified using predetermined thresholds.

#### Elevated Plus Maze (EPM)

The apparatus has a plus shaped maze with two open arms and two closed arms. Mice are placed in the center area for 10 minutes. The number of entries to each arm is recorded as well as time spent in there

#### Open Field (OF)

Each arena was 50 x 50 cm, with an opaque floor and illuminated at 25 lux. Mice were placed in the center of the arena and allowed to explore freely during 10 min. The floor was virtually divided into center (30 x 30 cm) and periphery. Time in each of those areas was recorded.

### 2.12. Cytokine measurement

Cytokine levels were measured using the MesoScale Discovery V-PLEX proinflammatory panel 1 (Meso Scale Diagnostics, Rockville, MD, USA) following manufacturer’s instructions. Samples and standards were incubated in the detection plate. After the plate was washed, detection antibodies were added. Signals were measured using the MSD instrument and concentrations were calculated based on the standard curves.

### 2.13. Immunohistochemistry (IHC)

Freshly isolated brains were sectioned around the mid area and fixed in formalin for 24 h. Then they were embedded in paraffin, sectioned at 4 µm thickness and baked overnight at 58°C.

For CD68: Sections were deparaffinized, hydrated and epitope retrieval was induced with Dako low pH target retrieval buffer (Agilent, Santa Clara, CA, USA) in a decloaking chamber (Biocare Medical, Pacheco, CA, USA) at 95°C for 20 min. Slides were washed and peroxidase activity was blocked with Dako quench reagent (Agilent). Anti-CD68 antibody (LS-C312677) was added for 1 h at room temperature followed by Dako envision anti-rabbit HRP polymer and Dako 3,3′-diaminobenzidine (DAB) (Agilent).

For 4HNE: The Ventana Discovery platform was used. Deparaffinization and antigen retrieval were performed on-board using CC2 mild conditions (Roche, Basel, Switzerland). Nonspecific staining was blocked with antibody block (Roche), then primary antibody against 4HNE was added (ab46545) and peroxidase activity was quenched. Slides were then incubated with Ventana anti-rabbit-HQ (Roche) anti-HQ-HR (Roche) and Ventana Chromomap DAB (Roche).

All slides were counterstained with Mayer’s hematoxylin and bluing water

For TUNEL assay, the Apoptag Peroxidase In Situ Apoptosis detection kit (Millipore- Merck KGaA, Darmstadt, Germany) was used on 4 µm sections from formalin fixed paraffin embedded tissue. Slides were deparaffinized, hydrated followed by PK digestion and quenching of peroxidase activity (Agilent). Equilibration buffer was applied and working TdT enzyme (prepared following manufacturer instruction) was added. Stop buffer was added, slides were washed and incubated with anti-DIG-HRP and DAB (Agilent). Slides were counterstained with Mayer’s hematoxylin and bluing water. Slides were dehydrated, cleared and mounted.

### 2.14. Cell culture

Cells were grown in a humidified incubator at 37 °C and 5% CO2. Human GBM LN18 cells were cultured in RPMI-1640 supplemented with 10% fetal bovine serum (FBS), 1% penicillin-streptomycin (P/S), 1% sodium pyruvate, 1% L-Glutamine solution and 1% HEPES solution. Human microglial HMC3 cells were cultured in EMEM (Corning) supplemented with 10% FBS and 1% pen-strep. Cells were purchased and were authenticated using Short Tandem Repeat by ATCC. Murine GBM GL261 cells were cultured in DMEM supplemented with 1% sodium pyruvate, 1% L-Glutamine solution and 10% FBS. Cells at passages below 10 were used in all experiments. GBM patient derived xenografts (PDXs) were obtained from the Mayo Clinic. G44 and G84 cells were cultured in DMEM supplemented with 10% FBS and 1% P/S. All reagents were purchased from Sigma-Aldrich unless specified otherwise.

For EV collection, 5x10^6^ cells were seeded in 15 cm petri dishes. When 80-90% confluent, cells were rinsed twice with PBS and fed with medium supplemented with EV-free FBS (System Biosciences) before receiving 6 Gy radiation treatment. Then, cells were incubated for 48 h and media was collected for EV isolation.

For microglial analyses, 3x10^5^ HMC3 cells were seeded in 6 well plates. 24 h later, cells were treated at 300 EVs/cell and downstream evaluations were performed 24h after treatment. Viability was assessed in cell suspension, using trypan blue and the Countess 3 FL Automated Cell Counter (Thermo Fisher Scientific).

### 2.15. Confocal microscopy

For internalization study, EVs were collected from LN18-RFP, a cell line transfected with RFP in the cell membrane. This allows for intrinsic fluorescence and avoids nonspecific staining. HMC3 cells were stained with green CellMask (Thermo Fisher Scientific) for the plasma membrane and Hoechst (Invitrogen-Thermo Fisher Scientific) for nuclear staining. Fluorescent EVs were added to HMC3 cells and confocal images were acquired with a Nikon AXR confocal microscope. Time course imaging was done acquiring a complete z-stack every 90 s. Images were acquired with sequential scanning to avoid bleed-through between channels

### 2.16. Image analysis using IMARIS

Image analysis for internalization experiments was performed using Imaris (Oxford Instruments Buckinghamshire, UK) to reconstruct z-stacks into 3D images. The surface detection function was used to identify cell boundaries and reconstruct cell volume. The surface detection function allowed identification of individual EVs. Parameters such as spot diameter, intensity threshold, background subtraction and distance to the surface were optimized to ensure accurate positioning in relation to the cell (surface), total count and minimize false positives.

### 2.17. Measurement of H_2_O_2_

Cells were seeded in 96 well plates at 30,000 cells/well. Cells were treated at 300 EVs/cell. 24 h after media was collected. H_2_O_2_ was measured using the Amplex Red Kit (Thermo Fisher Scientific) following manufacturer’s recommendations. Polyethylene glycol (PEG) - Catalase(PEG-Cat) was added to the cells as a negative control at a concentration of 500 U/µL, 24 h before measurement. Briefly, the Amplex Red reagent stock solution is combined with horseradish peroxidase (HRP) and reaction buffer. This working solution is mixed 1:1 with samples and incubated for 3h protected from light. Fluorescence was measured with a microplate reader (excitation 540 nm, emission 590 nm).

### 2.18. Metabolic measurements (mitochondrial and glycolysis)

Metabolic parameters were measured using the seahorse XFe96 analyzer from Agilent as described in (Miller, Xu et al. 2022). Mitochondrial function was measured using the Cell Mito Stress Test (Agilent) where sequential injection of electron transport chain inhibitors allows for determination of key parameters. Glycolytic function was assessed with the Glycolysis Stress Test (Agilent) where cells are forced to shift energy production to glycolysis followed by pharmacological inhibition to quantify glycolytic parameters.

### 2.19. P50 siRNA-mediated knockdown

Targeted gene knockdown of p50 was done using small interfering RNA (p50 siRNA) (cat. 4392420, Thermo Fisher Scientific) and the Stealth siRNA transfection protocol from Thermo Fisher Scientific. HMC3 cells were seeded in 6 well plates without antibiotics. When confluence reached 50%, cells were transfected with either p50-specific siRNA or scramble control siRNA using a mix of the oligomer and lipofectamine. Cells were incubated overnight and then media was changed to normal growth media. Knockdown efficiency was assessed 96 h post-transfection by Jess automated western blot confirming over 50% reduction of p50 compared to control cells.

### 2.20. Statistical Analysis

Data presented as mean ± SEM unless stated otherwise. Comparisons between two groups were performed using unpaired t-test. Comparisons between three or more groups were performed using one-way or two-way ANOVA followed by post hoc corrections as necessary. A p-value <0.05 was considered statistically significant. ∗P value < 0.05; ∗∗P value < 0.01; ∗∗∗P value < 0.001; ∗∗∗∗P value < 0.0001. All analyses were conducted using GraphPad Prism 11.

For proteomics the data missing valid accession IDs and protein without all the abundance values were excluded. For remaining proteins, missing values were imputed with the minimum abundance value for each one divided by √2. The resulting completed data was log2 transformed before all the downstream analysis.

Differential expression analysis was performed using the limma package (v3.65.3) (Ritchie, Phipson et al. 2015). Statistical significance was determined based on a false discovery rate (FDR) < 0.05, with p-values adjusted for multiple testing using the Benjamini–Hochberg (BH) method.

Gene set enrichment analysis (GSEA) was conducted using GSEA software (v4.4.0) (Subramanian, Tamayo et al. 2005, Subramanian, Kuehn et al. 2007) on ranked protein lists derived from DE analysis. Significant enriched pathways were identified based on FDR < 0.25.

GO enrichment analysis was performed using clusterProfiler (v4.17.0) (Yu, Wang et al. 2012) with BH method for multiple testing adjustment, and significant enrichment was defined using an FDR cutoff of 0.25.

## 3. RESULTS

### 3.1. CIRCULATING EXTRACELLULAR VESICLES ARE ELEVATED IN PATIENTS WITH GLIOBLASTOMA

To assess whether the concentration of EVs in circulation (also known as vesiclemia) is altered in glioblastoma, we quantified circulating EVs in serum from a small cohort of GBM patients (n=4) and matched healthy controls (n=5). The GBM cohort consisted of 4 patients who provided serum for analysis. The mean age was 59 years (range: [40-73]) with two males (50%) and two females (50%). Most patients (3/4, 75%) had their primary tumor in the temporal lobe with the remaining patient (1/4, 25%) having an overlapping lesion. Vesiclemia was significantly elevated in GBM patients compared to controls, while the overall EV size distribution remained similar between groups (Figure 1A-B). Transmission electron microscopy (TEM) further confirmed the presence of particles with EV morphology in the serum isolates (Figure 1C).

**Figure 1:**
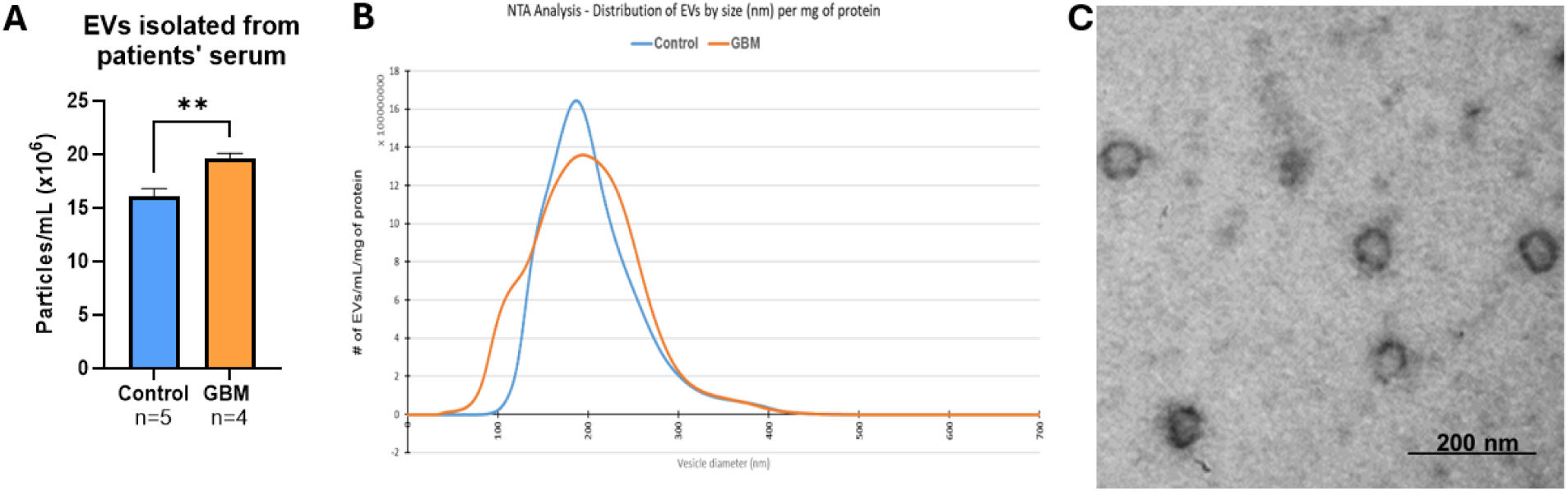
Vesiclemia is elevated in GBM patients. Circulating EVs were quantified in serum from GBM patients and healthy controls. **(A)** Total EV concentration measured by nanoparticle tracking analysis (NTA). GBM patients exhibited significantly higher EV levels compared to controls **(B)** Size distribution of EVs shows similar profile between groups. **(C)** Representative TEM image confirming the characteristic vesicular morphology of isolated EVs. ∗∗P value < 0.01.

These findings are consistent with previous reports demonstrating increased levels of circulating EVs in GBM patients (Ricklefs, Wollmann et al. 2024), confirming this observation in a different cohort and strengthening the relevance of our findings. Given the observed increase in systemic EVs in GBM patients, we next asked whether these vesicles have a role in GBM-associated cognitive dysfunction.

### 3.2. CHARACTERIZATION OF GBM-DERIVED EVs

To investigate the potential biological role of GBM-derived EVs we focused on two EV subtypes: EVs isolated from GBM cells (NON-RT-EVs) and EVs isolated from GBM cells that had been exposed to 6 Gy of ionizing radiation (RT-EVs), as most GBM patients receive radiation treatment.

Both EV subtypes were characterized using nanoparticle tracking analysis (NTA), transmission electron microscopy (TEM) and the presence of EV-specific markers. EV size and distribution were assessed across multiple cell lines and patient-derived xenografts (PDX). NON-RT-EVs and RT-EVs showed similar size distributions across all cell lines (∼200 nm, Figure 2A) and classical vesicular morphology (Figure 2D), indicating that any subsequent differences in EV abundance or specific markers, were not dependent on the physical properties of the vesicle. Radiation exposure also increased the total number of EVs released by GBM cells and PDXs (Figure 2B-C). Protein quantitation using Jess Automated Western blot confirmed enrichment of canonical EV markers CD9 and Flippase (Figure 2E).

**Figure 2:**
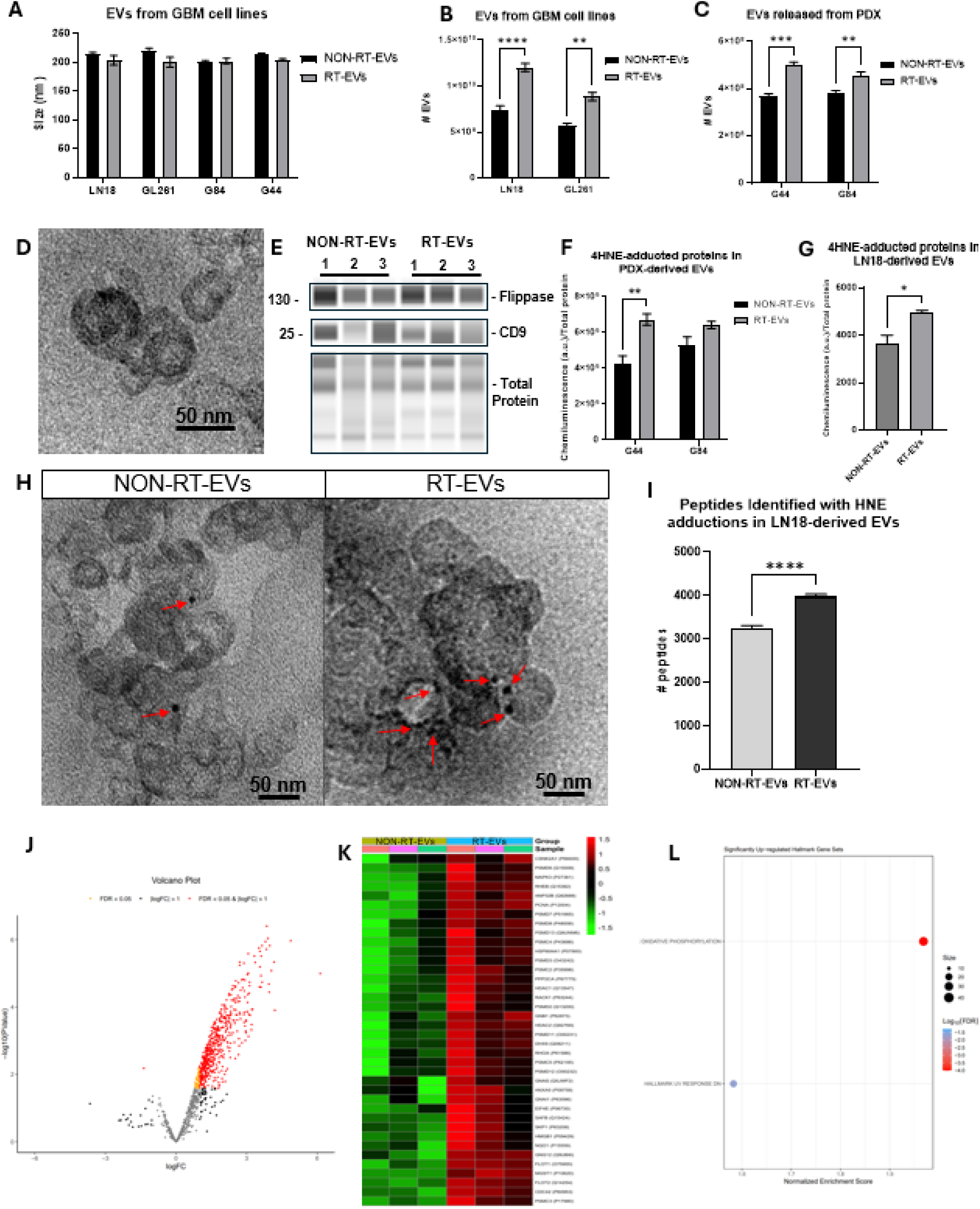
Characterization of GBM-derived EVs: NON-RT-EVs and RT-EVs were isolated from cell lines and PDX models. RT-EVs were isolated after cell exposure to 6 Gy. **(A-C)** NTA shows size distributions and concentration of NON-RT-EVs and RT-EVs across multiple cell lines and PDXs. Radiation treatment increased total EV release (RT-EVs>NON-RT-EVs). **(D)** Representative TEM confirms classical vesicular morphology. **(E)** Jess analysis demonstrates enrichment of canonical EV markers (CD…) and additional markers of interest. 4-hydroxynonenal (4HNE) content was measured by **(E)** immunogold labeling (representative image; red arrows indicate gold beads), **(F-G)** Jess analysis, and **(H)** peptide identification via mass spectrometry. Oxidative modifications were increased in RT-EVs in all methods. Proteomics analysis revealed enrichment of proteins in RT-EVs relative to NON-RT-EVs. **(I)** Heatmap showing over 500 significant proteins, clustered hierarchically (NON-RT-EVs in green, RT-EVs in orange. N=3 **(J)** Volcano plot of differentially expressed proteins (p<0.05 and fold change>1.5) in red and nonsignificant proteins in black. **(K)** Pathway analysis of significantly enriched proteins showing upregulation of oxidative phosphorylation and UV response hallmark gene sets. ∗P value < 0.05; ∗∗P value < 0.01; ∗∗∗P value < 0.001; ∗∗∗∗P value < 0.0001.

To assess oxidative stress associated with radiation, the levels of protein-bound 4-hydroxynonenal (4HNE) were measured. Immunogold labeling followed by TEM revealed increased 4HNE content in RT-EVs compared to NON-RT-EVs, with red arrows pointing at the gold beads marking 4HNE adducts (Figure 2H). Consistently, Jess analysis showed a quantitative increase in the 4HNE adducts present in RT-EVs (Figure 2F-G). To further validate the presence of 4HNE modifications, mass spectrometry analysis was performed to identify peptides bearing 4HNE adducts (Figure 2I), which confirmed the increase in 4HNE content is not only in the EVs as a whole but also in the proteins within the cargo. Overall, this confirms the alteration of redox-related cargo after radiation, both quantitatively and spatially.

Proteomic profiling of both EV subtypes was performed to identify molecular differences associated with exposure to radiation. Quantitative proteomic analysis identified over 500 significant proteins between both EV subtypes. Relative to NON-RT-EVs, RT-EVs show enrichment of numerous proteins as shown on the volcano plot analysis (Figure 2J), reflecting global remodeling of metabolic pathways rather than individual proteins. The heatmap highlights proteins involved with oxidative damage response, oxidative stress, cytokine release and the activation of NFκB pathway (Figure 2K). Consistently, pathway analysis highlighted the upregulation of the oxidative phosphorylation and UV response hallmark gene sets in RT-EVs (Figure 2L) indicating proteomic adaptations to radiation response.

These findings reveal the molecular differences in protein cargo between NON-RT-EVs and RT-EVs, with RT-EVs representing a subtype altered by radiation exposure that is released more from the cells, enriched in oxidative stress markers and has proteomic signatures consistent with the radiation that was received (6 Gy), suggesting that this treatment significantly altered energy metabolism and redox homeostasis. This served as the foundation for *in vivo* studies examining the biological impacts of both EV subtypes.

### 3.3. DIFFERENTIAL BIOLOGICAL EFFECTS OF NON-RT-EVs AND RT-EVs FOLLOWING INTRANASAL DELIVERY in vivo

To assess *in vivo* effects of GBM-derived EVs, female mice were treated with NON-RT-EVs or RT-EVs from GL261 cells using intranasal administration. Delivery to the brain was confirmed using fluorescently labeled EVs and *in vivo* fluorescence imaging (Supplementary figure 1). Two weeks later, behavioral testing revealed different significant effects between mice treated with NON-RT-EVs and RT-EVs.

Novel Object Recognition task was performed. The ratio of exploratory time directed toward the novel vs familiar object did not differ between groups (p>0.05) indicating comparable recognition performance (Figure 3A). However, animals treated with RT-EVs required significantly more time to reach 20 s of cumulative exploration, indicating delayed object exploration despite preserved total exploration time. (Figure 3B). This suggests a difference in the dynamics of object-directed exploration and processing of novel stimuli rather than a reduction in exploratory activity. Importantly, all groups had comparable times of total exploration (Figure 3C). Consistent with this interpretation, locomotor activity and preference for periphery vs center in the open field (Figure 3D-E) and performance in the elevated plus maze (Figure 3F) were similar between groups, arguing against locomotor or anxiety related explanations for the cognitive delay in object exploration.

**Figure 3:**
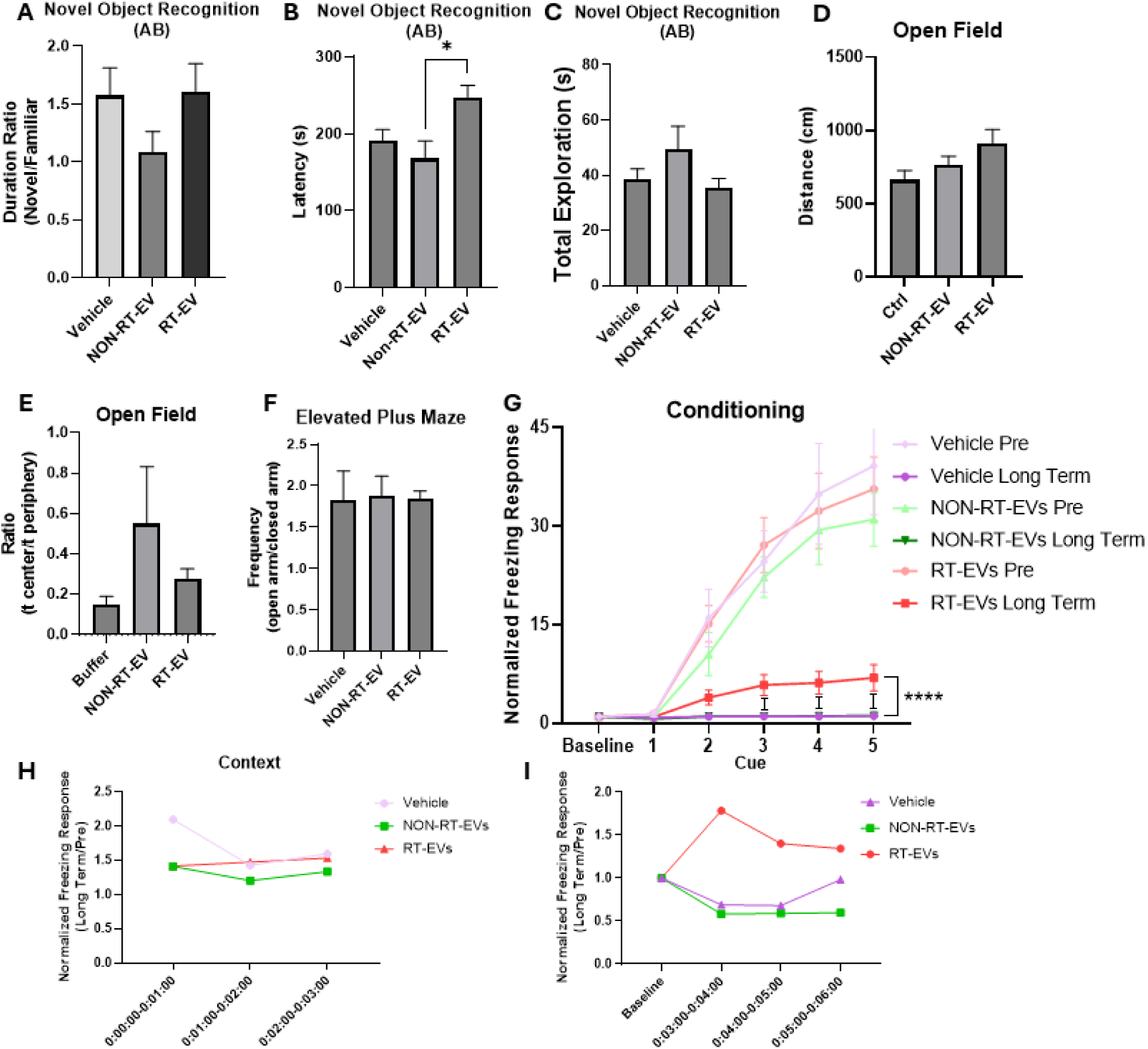
RT-EVs cause cognitive alterations following intranasal delivery. (A-C) Novel Object recognition test shows comparable recognition and exploration across groups but significant longer time to complete the task in RT-EV-treated mice. Open field test across groups displaying comparable (D) total distances traveled and (E) ratios of time spent in the center vs periphery, indicating comparable locomotion and anxiety-like behaviors respectively. (F) Elevated plus maze showing the relation of time spent in the open vs closed arm as confirmation of absence of anxious behaviors. Fear conditioning test presenting (G) freezing response to conditioning conditions before and after EV treatment, (H) context results and (I) cue response. Data presented as the ratio of averaged freezing behavior long term vs before treatment (I) Total distance traveled in the open field test demonstrates comparable locomotor activity across groups. N=10/group. ∗P value < 0.05; ∗∗∗∗P value < 0.0001.

Animals treated with RT-EVs also showed impairments in cognitive performance related to stress response. Fear conditioning was performed before EV treatment, where all groups showed comparable freezing response across timepoints, indicating intact initial learning. A month later, when the animals were rechallenged with the conditioning protocol, mice treated with RT-EVs displayed significantly different conditioning behavior, whereas controls and NON-RT-EV-treated mice showed little to no freezing (Figure 3G). The context portion showed similar results across all treatment groups, suggesting that contextual memory is not affected (Figure 3H). Consistently with the conditioning results, when mice were exposed to the cue portion of the test after a month, mice treated with RT-EVs showed an enhanced sensitivity to stimuli, showing that cued memory is impaired long term (Figure 3I). In general, both behavioral assays show neurobiological deleterious effects of RT-EVs. The NOR shows mild acute deficiency as seen on the latency results and the FC shows long term changes in memory and learning, specifically in cued memory but not contextual. Importantly, baseline locomotor activity was comparable across all groups after EV administration as shown by the distance traveled in the open field test (Figure 3D), demonstrating that treatment did not affect locomotion and differences in freezing behavior are not due to altered mobility.

In addition to behavioral changes, biochemical analyses were performed on brains harvested from these mice. Jess Automated Western Blot showed that levels of 4HNE adducts were increased in neurons isolated from brains treated with either EV subtype (Figure 4A), suggesting that cellular responses are being triggered by GBM-derived EVs.

**Figure 4:**
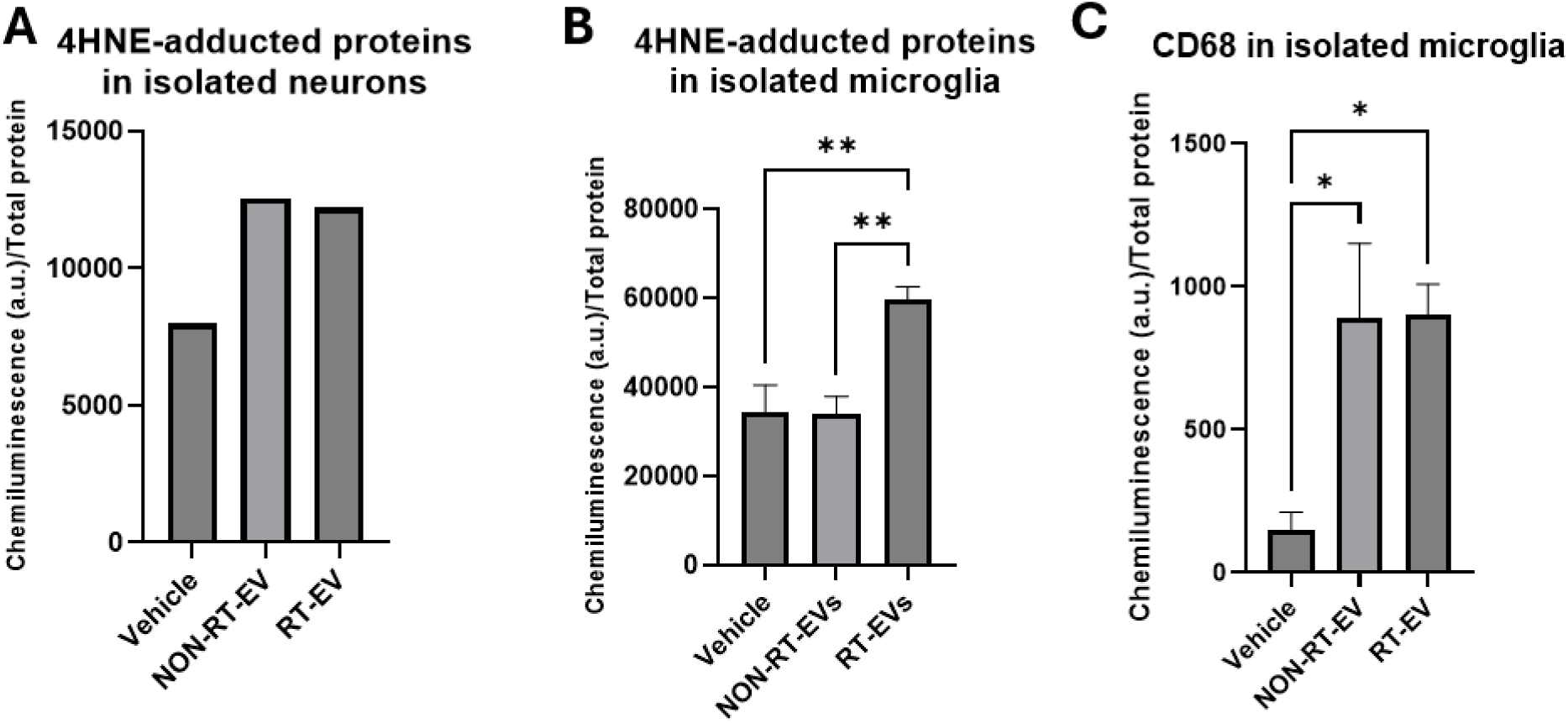
RT-EVs induce biochemical changes in the female brain following intranasal delivery. (A) Neurons isolated from mice treated with either NON-RT-EVs or RT-EVs show increased 4HNE adducts. Data presented as a single measurement of 5 mice per group (B) Microglia isolated from mice treated with RT-EVs show increased 4HNE adducts (C) CD68 expression is significantly increased in mice treated with either NON-RT-EVs or RT-EVs. N=5/group. ∗P value < 0.05; ∗∗P value < 0.01.

Additionally, microglia cells were isolated. Jess Automated Western Blot showed increased levels of 4HNE adducts in microglia isolated from brains of mice treated with RT-EVs (Figure 4B), which is consistent with the findings in neurons and suggests that the oxidative stress spreads to different cells in the brain, Additionally, the levels of CD68 were elevated in both groups of treated mice (Figure 4C), suggesting that either EV type can trigger microglial activation

Together, these findings indicate that RT-EVs exert stronger biological effects *in vivo* compared to NON-RT-EVs after intranasal delivery. Given that radiation therapy is part of the standard of care for GBM, RT-EVs represent a clinically relevant EV population to be studied further. Additionally, intranasal delivery of EVs provided limited control over the precise quantity of EVs being delivered to the brain. Therefore, RT-EVs were selected for further investigation using intracranial delivery, which allows for confirmation of brain delivery and precise control of EV dosing.

### 3.4. INTRACRANIAL DELIVERY OF RT-EVs REVEALS COGNITIVE IMPAIRMENT IN FEMALE MICE/IN VIVO AND INCREASED OXIDATIVE STRESS

To further determine the functional effects of RT-EVs, we evaluated the impact of RT-EVs delivery using intracranial injections to provide more controlled brain exposure. EVs were delivered to the somatosensory cortex to avoid major structures of the brain. There was no significant difference in mice weights between treatment groups. Ten days after administration, animals were subjected to behavioral assays to assess cognitive function.

In the open field test, animals traveled similar distances (Figure 5A) and displayed comparable ratios of time in the center vs the periphery (Figure 5B), indicating that locomotion was not affected by treatment and anxiety response is comparable between groups. Consistently, performance in the light-dark box test did not differ between groups, as measured by duration in the light area and distance traveled (Figure 5C), confirming equivalent anxiety-like behaviors. However, qualitative analysis of exploratory patterns within the light compartment revealed different spatial patterns between groups (Figure 5D), showing differences in exploratory strategy. These suggest that subsequent behavioral differences are not attributable to alterations in locomotion or basal anxiety levels.

**Figure 5:**
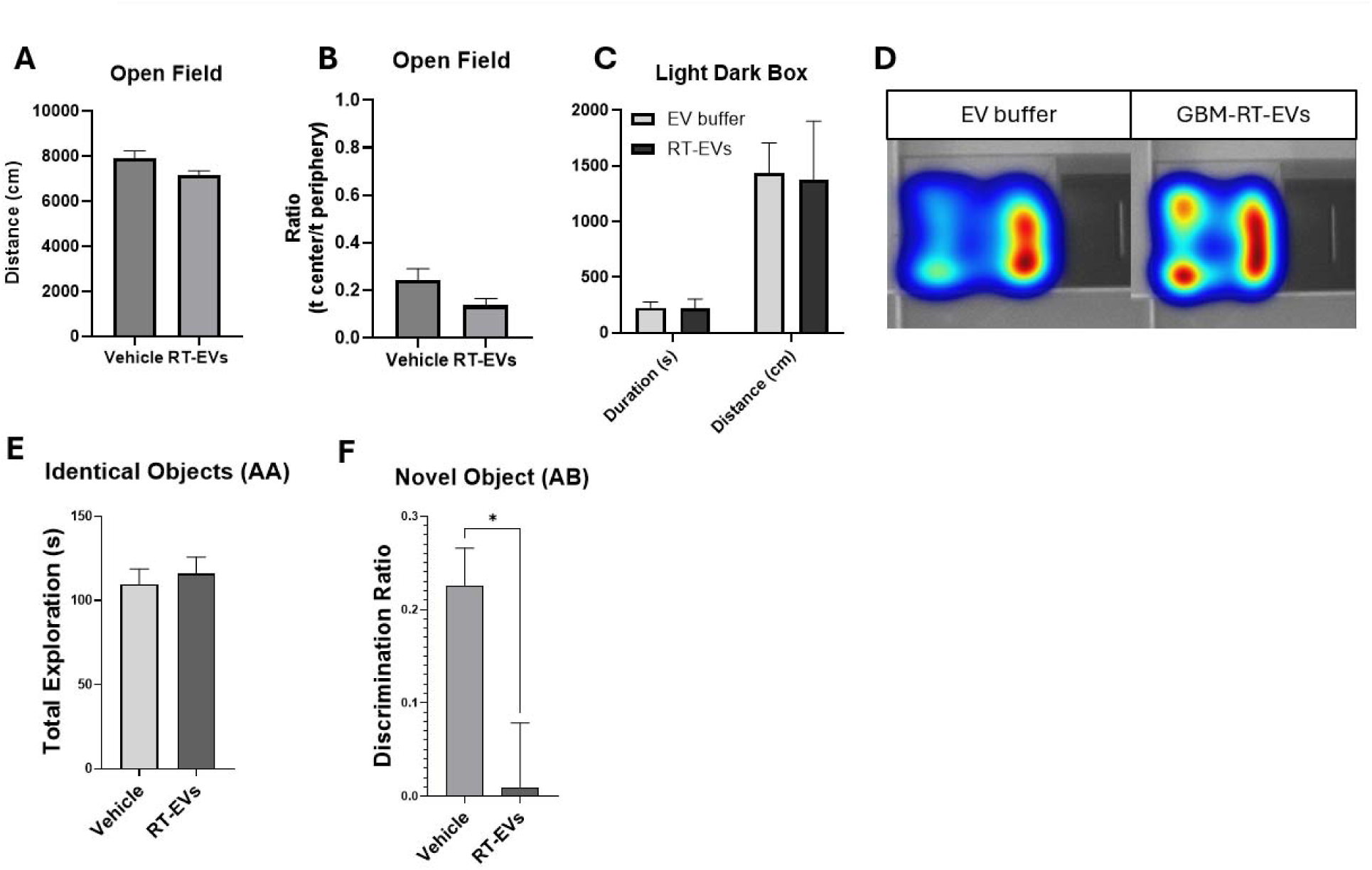
Intracranial delivery of GBM-RT-EVs induces cognitive deficits in female mice. Performance in the open field **(A)** Total distance traveled and **(B)** ratio of time spent in the center vs the periphery of the arena. Performance in the light-dark box showing **(C)** duration and distance spent in the compartments and **(D)** representative heatmaps of exploration, highlighting differential exploration between groups. Performance in the Novel Object Recognition test, showing **(E)** total exploration during the acclimation period and **(F)** the discrimination index during the test phase, indicating reduced recognition memory in mice treated with RT-EVs, while locomotive and exploratory activity remained similar. N=4/group for OF and light dark box. N=9/group for NOR. ∗P value < 0.05.

In the NOR task, RT-EV-treated mice exhibited a significant reduction in discrimination index compared with control animals, indicating impaired recognition memory (Figure 5F). Total exploration time during AA phase did not differ between groups (Figure 5E), indicating comparable baseline exploration and locomotive capabilities, which confirms that the observed deficit was not attributable to altered exploratory behavior.

These findings reinforce the behavioral effects observed with intranasal delivery and demonstrate that RT-EVs can induce measurable cognitive deficits *in vivo*.

Subsequently, animals were euthanized and tissues were collected for analysis. TUNEL assay revealed increased TUNEL-positive cells in the brains of mice treated with RT-EVs (Figure 6A-B), suggesting an increase in apoptotic cells. Immunohistochemical analysis revealed increased 4HNE adducts in the hippocampal area of mice treated with RT-EVs (Figure 6C-D). Jess analysis revealed an increase of 4HNE adducted proteins (Figure 6E-F) and NOX4 (Figure 6G-H) in brain lysates of mice treated with RT-EVs, suggesting enhanced oxidative stress and lipid peroxidation. In contrast, markers associated with neuronal health like DCX, NeuN and Neurofilament were decreased compared to controls (Figure 6I-J), indicating neuronal damage.

**Figure 6:**
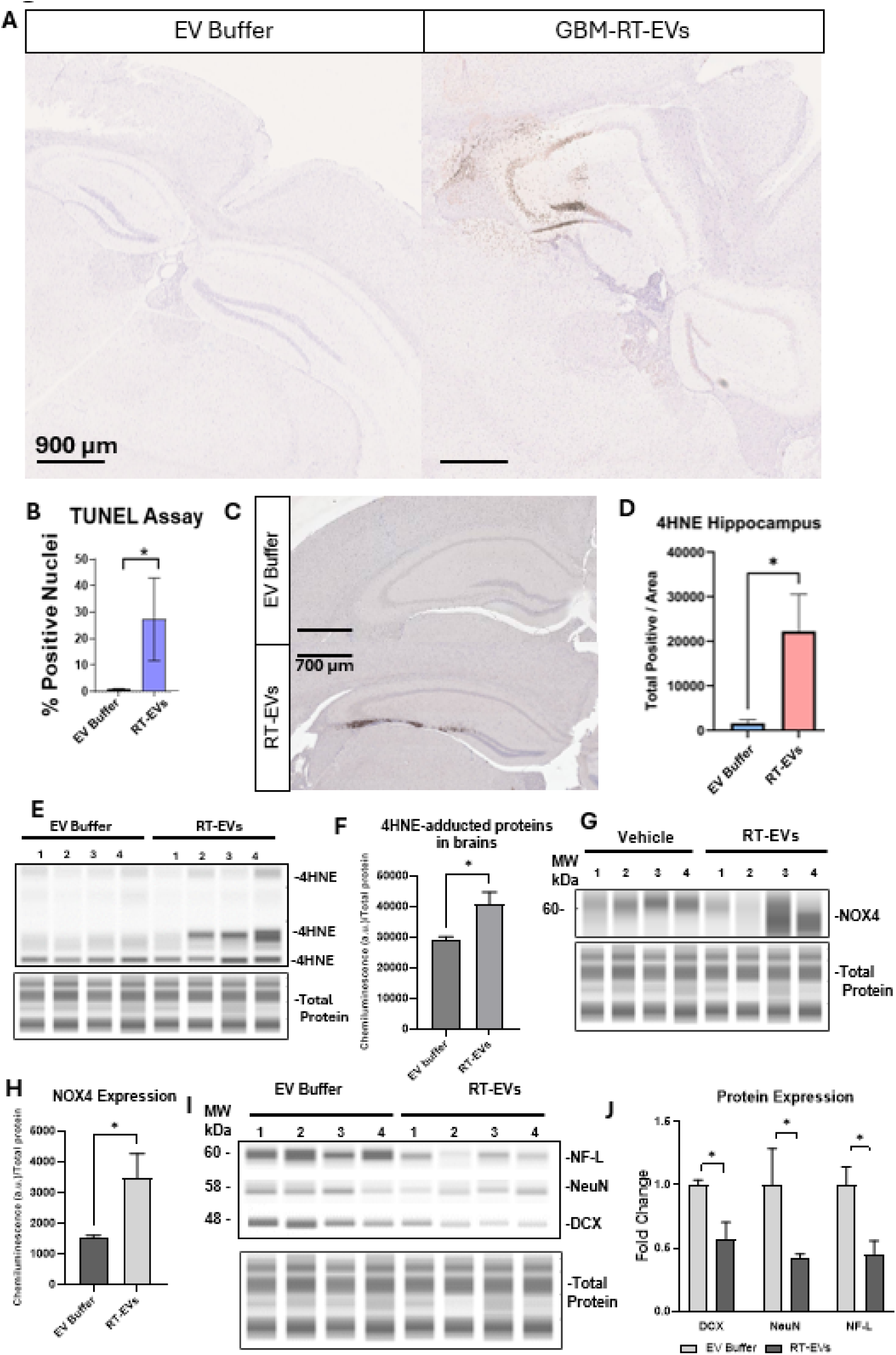
GBM-RT-EVs cause molecular alterations in the female brain upon intracranial administration. **(A)** Representative images of TUNEL staining and **(B)** quantification of TUNEL-positive nuclei. (C-D)-. Representative blot images and quantification of **(E-F)** 4HNE-adducted proteins, **(G-H)** NOX4 and **(I-J)** neuronal markers including DCX, NeuN and Neurofilament. ∗P value < 0.05.

Given the evidence of oxidative modifications within RT-EVs, the increase in oxidative stress and neuronal damage markers in the brains of animals treated with RT-EVs and the established link between oxidative stress, neuroinflammation and cognitive dysfunction, we next investigated the involvement of NFκB signaling following RT-EVs exposure

### 3.5. RT-EVs INCREASE P50 LEVELS IN THE FEMALE BRAIN

To investigate potential molecular mechanisms underlying the cognitive phenotype following RT-EV treatment, we examined activation of inflammatory pathways in the brain given the increase in CD68 expression observed in microglia following EV administration in section 3. We focused on the involvement of NFκB pathway, a key regulator of inflammatory and redox responses in the brain. Among the NFκB family, the p50 subunit is associated with canonical activation and has been implicated in neuroinflammatory processes.

We first assessed the p50 levels in microglia isolated from the brains of mice treated intranasally as described in section 3. Jess analysis revealed increased p50 levels in microglia isolated from animals treated with both NON-RT-EVs and RT-EVs (Figure 7A-B). Notably, the increase was more pronounced in microglia derived from RT-EV-treated mice, consistent with the previously described enhanced biological activity associated with RT-EVs.

**Figure 7:**
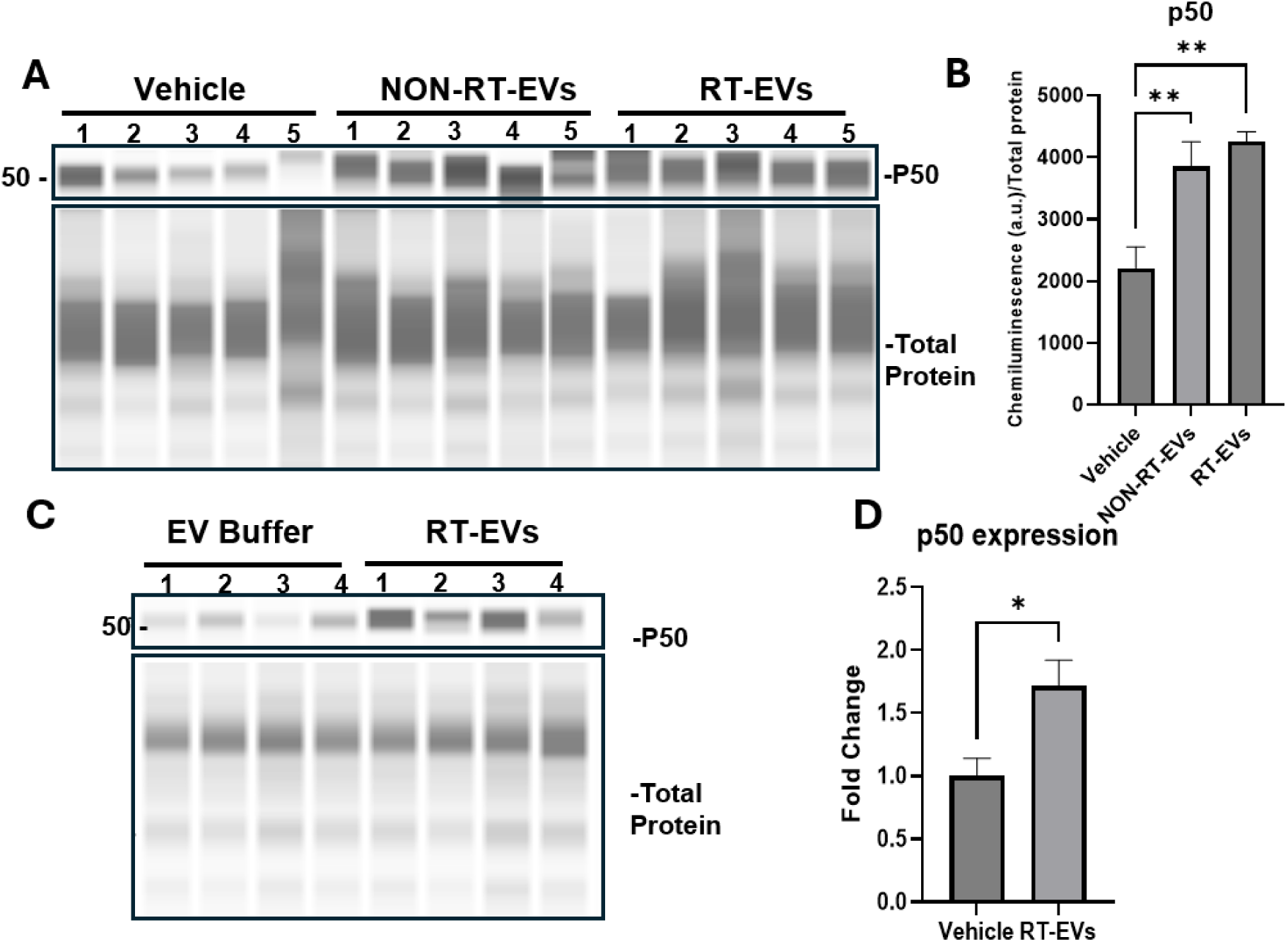
RT-EV exposure increases NFκB p50 levels. Jess analysis of p50 in microglia isolated from brains of mice treated with NON-RT-EVs or RT-EVs using intranasal delivery. (A) Representative blot and (B) quantification of area under the curve showing higher p50 levels in microglia from RT-EV-treated mice compared to those from NON-RT-EV treated mice and controls. Jess analysis of total brain lysates from mice treated with RT-EVs intracranially (C) representative blot and (D) quantification of area under the curve showing significant increase in p50 compared with vehicle controls. ∗P value < 0.05. ∗∗P value < 0.01.

To determine if this activation was evident at the tissue level, we next analyzed total brain lysates from mice receiving RT-EVs intracranially as described in section 4. Consistent with the microglial findings, mice treated with RT-EVs showed a significant increase in p50 protein levels when compared to control animals (Figure 7C-D)

Together, these results indicate that the exposure to GBM-derived RT-EVs is associated with microglial activation of NFκB-related signaling pathways in the brain. Given the established role of NFκB signaling in the regulation of microglia activation, including oxidative stress responses, we next examined changes in microglial phenotype.

### 3.6. RT-EVs INDUCE MICROGLIAL ACTIVATION IN VITRO

To investigate whether RT-EVs directly affect microglia in vitro, we exposed human microglia to RT-EVs and assessed vesicle internalization, activation phenotypes and redox response.

#### Internalization and morphology

Microglial cells were exposed to RT-EVs derived from RFP-LN18 (LN18 cells transfected to express RFP in their membrane) and monitored using confocal microscopy. The results confirmed punctate intracellular fluorescence within 5 minutes (Figure 8A-B). Images from z-stack were used for 3D reconstruction using IMARIS. Each EV was identified and tracked to determine its position. Internalization was confirmed using the position of each EV relative to the surface of the cell, indicating EV uptake rather than surface interactions (Figure 8C-D). Quantitative analysis demonstrated a time-dependent increase in the intensity inside the cells corresponding to EVs internalized (Figure 8E). EVs spread throughout the cell with no preferential accumulation at a specific site. Using light microscopy, we assessed microglial morphology after treatment. Images indicated that microglia treated with RT-EVs exhibited both amoeboid and hyper-ramified morphologies compared to control cells (Figure 8F). This is consistent with activation.

**Figure 8:**
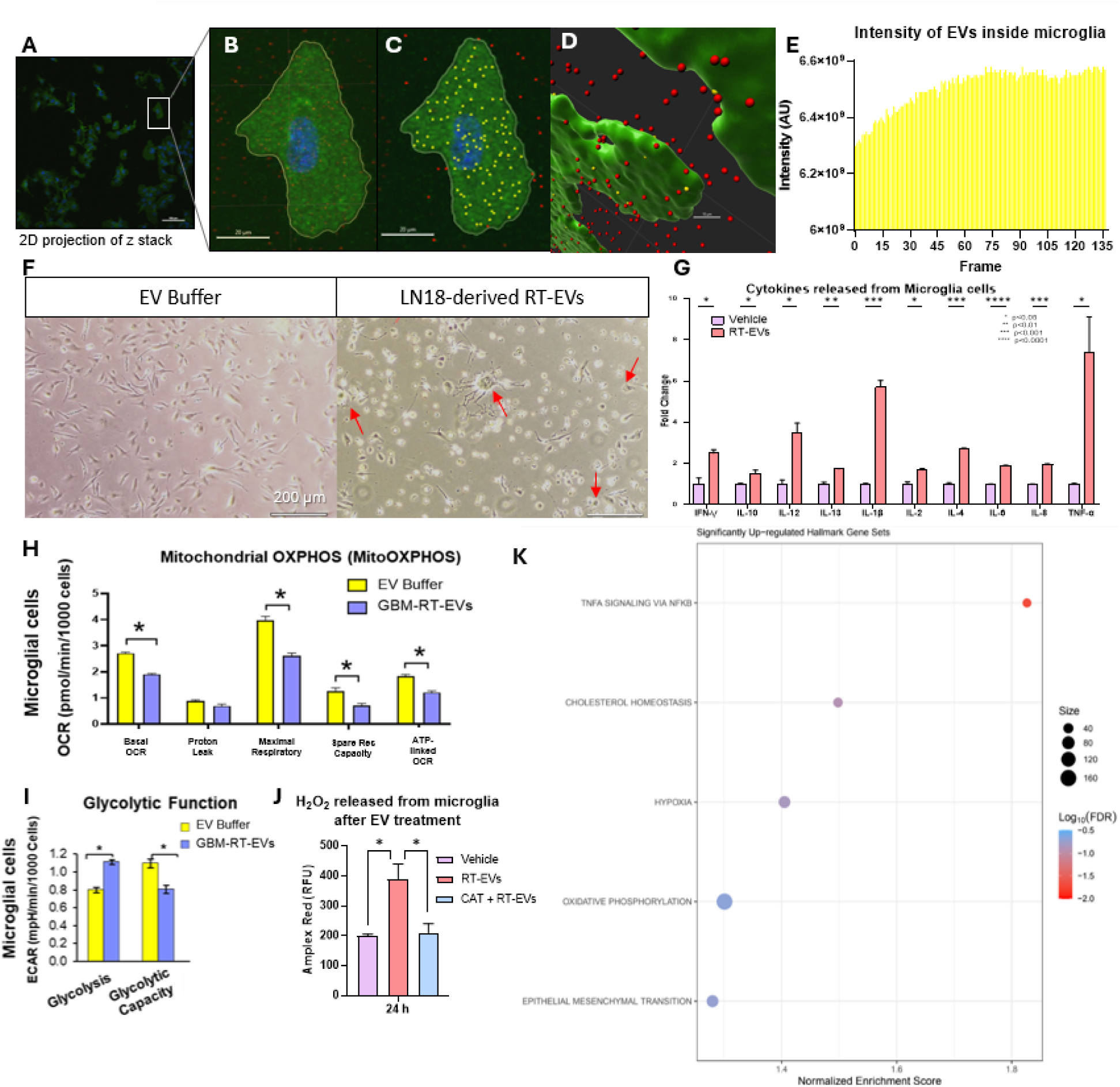
RT-EVs induce microglial activation in vitro: Confocal imaging and 3D reconstruction and analysis (A) Representative maximum intensity projection of z-stack showing cytoplasm (green), nuclei (blue) and RT-EVs (red) labeling. (B) Higher magnification of a representative microglia cell, illustrating RT-EVs internalization. (C) Individual EV identification using spot detection in IMARIS. EVs classified inside (yellow) and outside (red) the cell. (D) Representative 3D reconstruction of the cell generated from the confocal z-stack using IMARIS, illustrating spatial distribution of EV internalization. (E) Quantification of EV fluorescence intensity inside the representative cell, showing the time-dependent increase in internalization. (F) light microscopy images showing microglial morphology in control vs RT-EV-treated cells. Red arrows point to amoeboid and hyper-ramified cells. (G) Cytokine levels measured in culture media of microglial cells. Seahorse analysis showed (H) decreased mitochondrial respiration and (I) increased glycolytic activity (J) H_2_O_2_ released from microglial cells is increased upon RT-EVs treatment. Catalase abolishes the signal confirming H_2_O_2_ specificity. (K) Proteomic analysis of RT-EV-treated microglia showing upregulation of specific gene sets. ∗P value < 0.05; ∗∗P value < 0.01; ∗∗∗P value < 0.001; ∗∗∗∗P value < 0.0001.

#### Cytokine release

Following RT-EV administration to microglia, cytokine levels were quantified in culture media. RT-EV treatment significantly increased the secretion of the pro-inflammatory cytokines IFN-γ, IL-1β, IL-6, and TNF-α. Levels of IL-10 were also increased (Figure 8G). This indicates the activation of pro-inflammatory signaling that might be accompanied by regulatory feedback mechanisms.

#### Metabolic alterations

RT-EV exposure resulted in increased microglial glycolytic activity and a decrease in mitochondrial respiration (Figure 8H-I), suggesting a shift in metabolic state consistent with microglial activation.

#### ROS production

Microglia treated with RT-EVs produced significantly higher levels of hydrogen peroxide compared with controls, showing alterations in redox state. This increase was observed as early as 3 h (data not shown) and sustained at 24h (Figure 8J).

#### Proteomic profiling

Proteomic analysis of microglial lysates following RT-EV exposure revealed enrichment of proteins in the TNF-α signaling pathway via NFκB (Figure 8K) when compared to controls. Other pathways upregulated were hypoxia, oxidative phosphorylation and epithelial mesenchymal transition.

Together, these data demonstrate that RT-EVs are directly internalized by microglia and induce activation, including cytokine release, metabolic reprogramming, and H_2_O_2_ production. Results from proteomic profiling provide a mechanistic link to the indications of NFκB activation *in vivo* (section 5) and confirm NFκB involvement in the functional responses *in vitro*.

### 3.7. H_2_O_2_ FROM RT-EV-ACTIVATED MICROGLIA INDUCES NEUROTOXICITY IN VITRO

To determine if oxidative stress generated by RT-EV-activated microglia could compromise neuronal integrity, we established a co-culture setting of human neurons with microglia that was previously exposed to RT-EVs. Neuronal viability and morphology were assessed after 48 hours. Neurons co-cultured with microglia previously treated with RT-EVs exhibited a significant reduction in viability compared to neurons in co-culture with untreated microglia (Figure 9B). Morphological assessment revealed profound alterations, where neurons treated with RT-EVs lost their characteristic projections and showed widespread blebbing compared with control cells indicating neuronal injury (Figure 9A). Importantly, pretreatment with catalase, an enzyme that decomposes hydrogen peroxide, fully rescued neuronal viability and prevented morphological changes (Figure 9A-B) confirming that microglial H_2_O_2_ was responsible for the observed neuronal toxicity.

**Figure 9:**
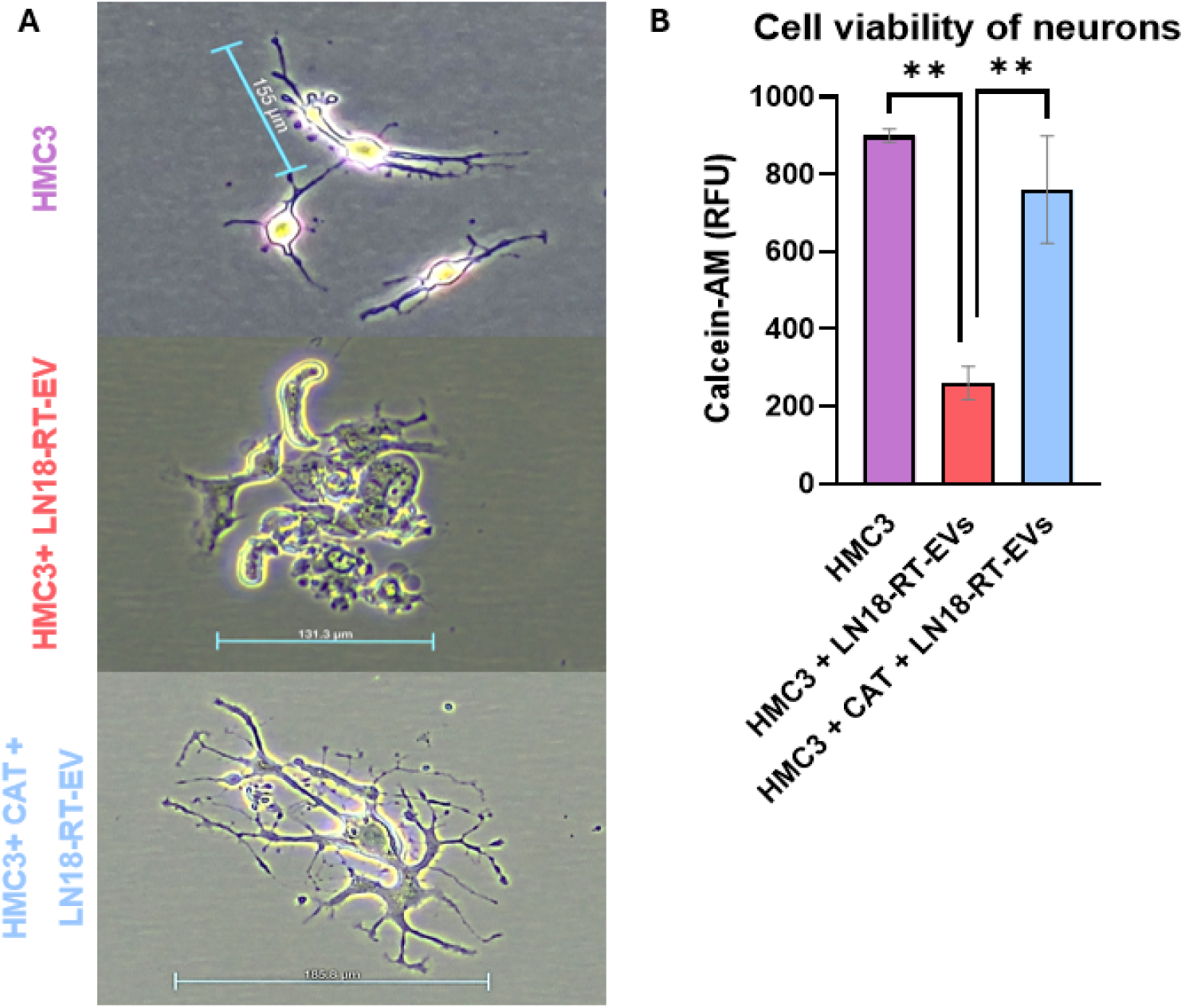
Microglial H_2_O_2_ produced upon RT-EV exposure causes neuronal injury: **(A)** Representative image of neuronal morphology and **(B)** quantification of neuronal viability. Catalase rescues neuronal survival and morphology confirming H_2_O_2_ dependent toxicity

This demonstrates that the oxidative stress induced by RT-EVs in microglia, is sufficient to damage neurons, providing a mechanistic link between microglial activation and ROS production as drivers of the cognitive phenotypes observed *in vivo* (sections 3-4).

### 3.8. H_2_O_2_ SECRETION INDUCED BY RT-EVs IN MICROGLIA IS NF**κ**B DEPENDENT

To assess if NFκB signaling is required for RT-EV-induced oxidative responses in microglia, we performed targeted KD of NFκB p50 in human microglia (HMC3) prior to exposure to RT-EVs (Figure 10A). Microglia with p50 KD did not show an increase in H_2_O_2_ release upon RT-EV exposure compared with control cells. In contrast, microglia treated with scrambled control siRNA had a significant increase in H_2_O_2_ secretion consistent with previous experiments (Figure 10B). Additionally, Jess analysis showed that RT-EVs increased 4HNE adduction and CD68 expression in control microglia cells (Figure 10C-D). Conversely, cells with p50 KD had no change in the 4HNE adduction or CD68 expression upon RT-EV treatment. In combination, this confirms that NFκB is required for RT-EV induced microglial activation and the production of microglial H_2_O_2_ upon RT-EVs exposure, which is neurotoxic.

**Figure 10:**
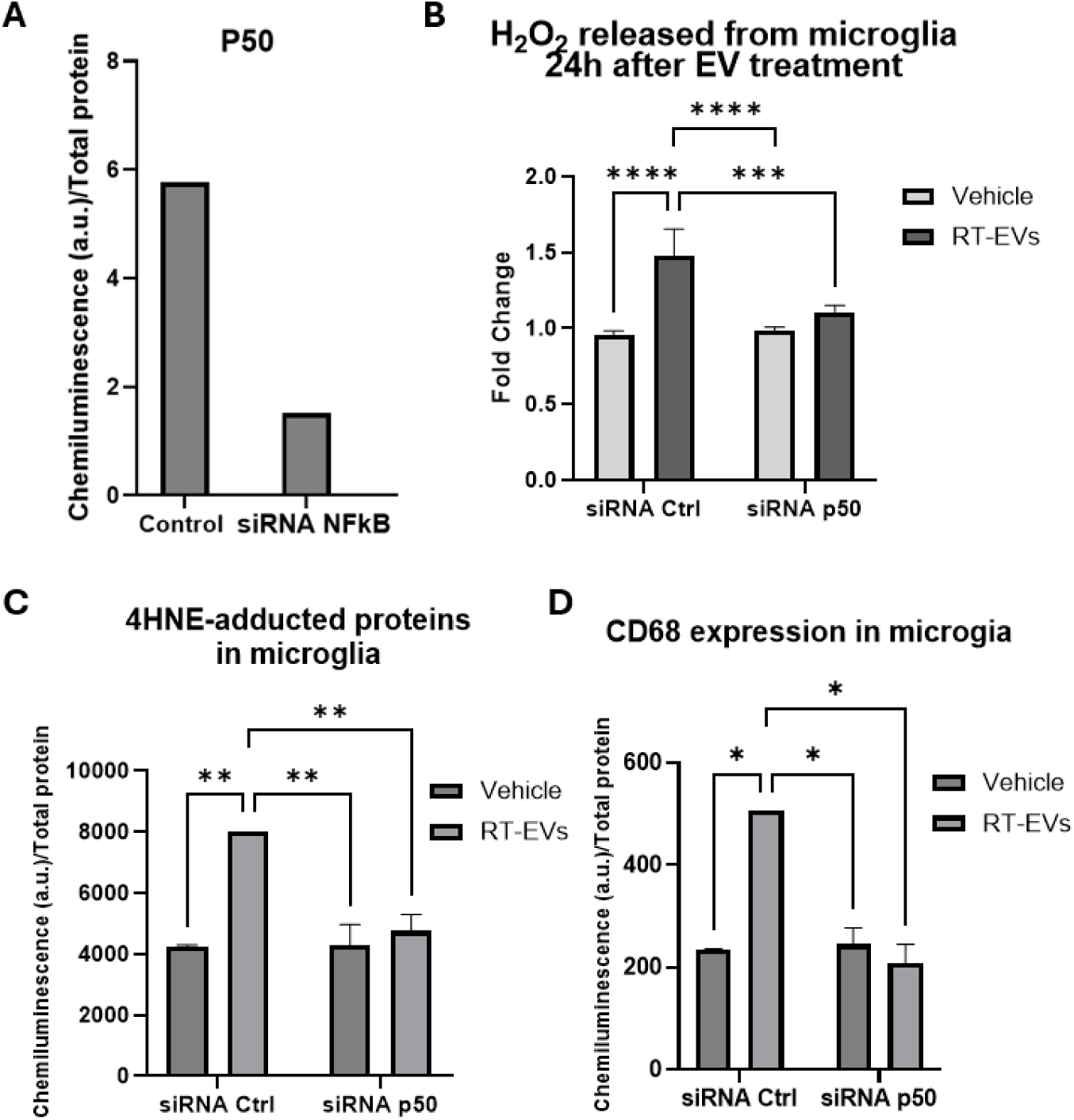
Reduced expression of NFκB p50 in microglia abrogated RT-EV-induced activation: (A) Confirmation of NFκB p50 reduced expression in microglia via Jess (B). Quantification of H_2_O_2_ production in microglia transfected with scrambled siRNA or p50-targeting siRNA followed by RT-EV exposure, showing abrogation of H_2_O_2_ increase in p50 KD cells. Jess analysis showing reduction of (C) 4HNE-adducted proteins and (D) CD68 expression upon p50 KD. ∗P value < 0.05; ∗∗P value < 0.01; ∗∗∗P value < 0.001; ∗∗∗∗P value < 0.0001

**Figure 11:**
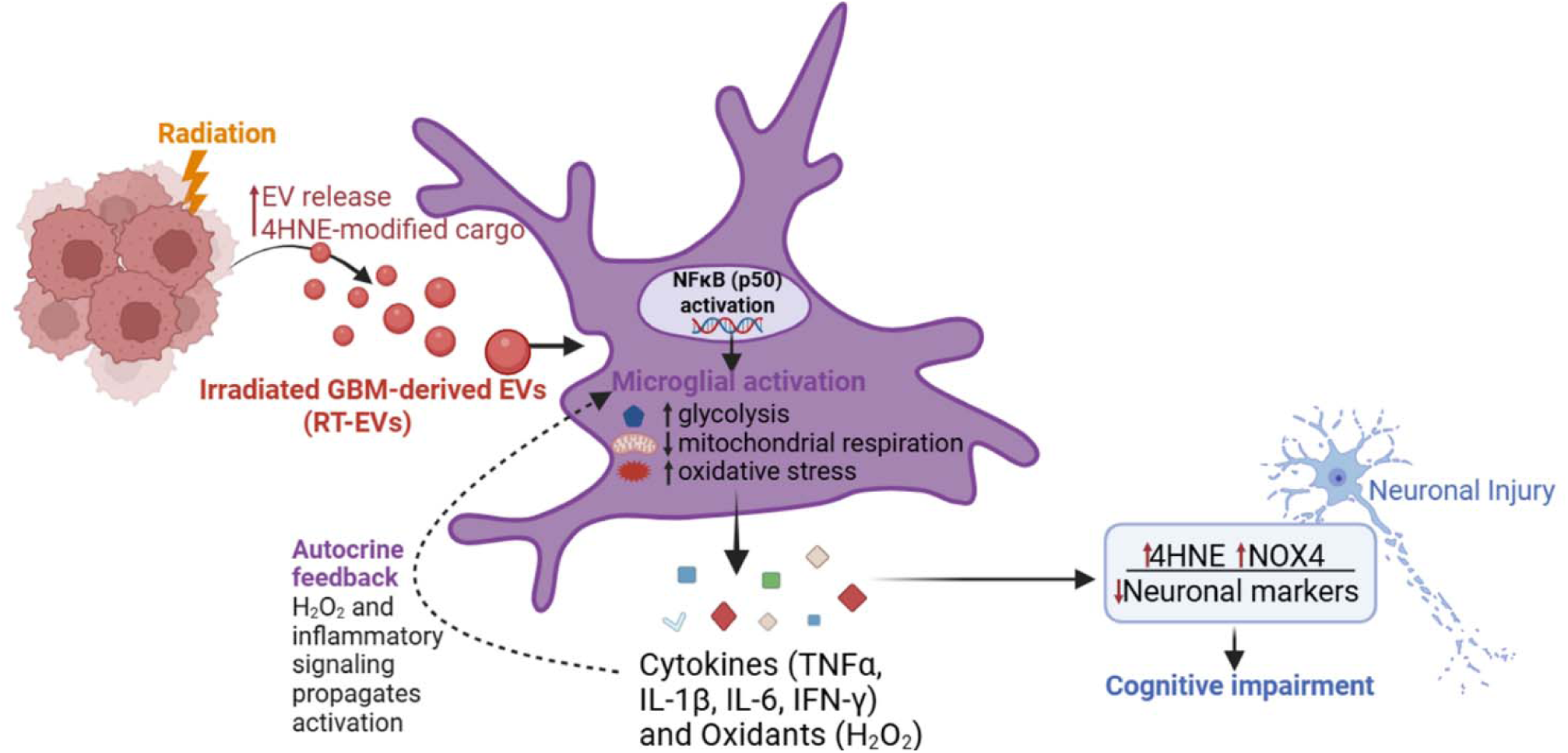
Proposed mechanism. Following radiation treatment, glioblastoma cells increase their EV release and the 4HNE modified cargo. RT-EVs activate microglia cells via NFκB triggering the transcription of oxidative and inflammatory factors, leading to secretion of cytokines and H2O2, which can directly interact with neurons and other cells, as well as return to the cell of origin to propagate the inflammatory response. Solid arrows indicate established pathways and dashed arrows represent indirect or proposed interactions

These findings show that NFκB activation is upstream of H_2_O_2_ production in microglia exposed to RT-EVs, providing a mechanistic explanation for the link between RT-EVs, microglial oxidative stress and neuronal dysfunction.

Collectively, this work established that RT-EVs cause cognitive decline *in vivo* in combination with increased oxidative damage and neuronal injuries. Additionally, RT-EVs cause microglial activation, triggering H_2_O_2_ production and neuronal death. Overall, our findings reveal a mechanistic pathway linking GBM-derived RT-EVs to cognitive dysfunction, providing a link between tumor-derived EVs, microglial redox signaling and brain dysfunction.

## 4. DISCUSSION

In this study, we demonstrate that EVs released by GBM cells after radiation are biologically distinct from EVs released under non-irradiated conditions and are sufficient to impair cognitive performance *in vivo*. Mechanistically, RT-EVs are internalized by microglia, activate NFκB-associated inflammatory signaling, promote metabolic reprogramming, and induce neurotoxic H_2_O_2_ production that compromises neuronal viability. These findings support a model in which radiation-altered tumor EVs act as systemic and local mediators of brain dysfunction in GBM.

Cognitive dysfunction is a major and clinically meaningful consequence of glioblastoma, whether it arises from the tumor itself and from its treatment. Defining the underlying mechanisms may improve our understanding of tumor-associated cognitive impairment and help identify therapeutic targets.

There are other factors that can contribute to cognitive alterations in GBM patients including the mass effect of the tumor (Prasanna, Mitra et al. 2019), direct radiation-induced tissue damage (Fu, Zhang et al. 2024) and chemotherapy (named chemobrain)(Rao, Bhushan et al. 2022). However, here we show that GBM-RT-EVs are sufficient to elicit redox imbalance in the brain, particularly in microglia cells via canonical activation of NFκB (via p50) followed by neuroinflammatory responses, expressed as cytokine release (IL-6, IL-1β, IL-10, TNFα), oxidative stress (H_2_O_2_ production, NOX4 and 4HNE increase), and subsequent cognitive impairment (NOR, FC, and others).

Extracellular vesicles have emerged as modulators of organ function and cancer progression (Urabe, Kosaka et al. 2020, Elias, Hadjiyiannis et al. 2025). Here, we demonstrate that GBM-RT-EVs cause measurable behavioral outcomes, neuroinflammatory responses and oxidative damage, significantly impacting brain homeostasis. Stress conditions like hypoxia and radiation exposure have been linked to an increase in EV release (Szatmari, Hargitai et al. 2019, Crewe 2023). Particularly, the H_2_O_2_ increase upon radiation therapy has been linked to an increase in EV release in a dose-dependent manner (Miller, Xu et al. 2022). This could explain the increase in EV numbers that we observed both in patients and *in vitro*, supporting the notion that radiation modulates EV release in glioblastoma. Additionally, evidence suggests that EVs derived under pathological conditions can alter the function of the CNS and influence behavioral outcomes (Wang, Li et al. 2022, Ma, Shin et al. 2023, Wu, Deng et al. 2025), supporting the notion that RT-EVs are sufficient to drive the cognitive alterations observed in this study.

Research shows that EVs can induce inflammatory responses in cells upon internalization, specifically, it’s been shown that EVs are sufficient to trigger NFκB activation in target cells (Papareddy, Tapken et al. 2024). Here, we show that RT-EVs modulate microglial activation via NFκB signaling, specifically the p50 subunit. Further studies should aim to investigate this activation in vivo, using murine conditional models of microglia-specific NFκB knockdown.

Despite these findings, and many other, pointing at EVs as inflammatory activators, the exact mechanism of how RT-EVs activate microglia remain to be understood. A possible explanation could involve the 4HNE content in these EVs. Previous studies (Chatterjee, Zhu et al. 2018) and our findings show how radiation increases 4HNE adducts in cells and EVs. Additionally, 4HNE on its own is implicated in microglial activation, induction of Nrf2, causing oxidative stress (Cumaoglu, Agkaya et al. 2019). Consequently, the high levels of 4HNE present in RT-EVs could be responsible for triggering the initial activation of NFκB in microglia cells.

Another possibility lies in the proteomic profile of RT-EVs. Although studies linking individual proteins to microglial activation are limited, alterations in overall protein content have been linked to microglial activation. We identified a global protein remodeling in RT-EVs, with upregulation of proteins related to oxidative damage response, oxidative stress, cytokine release and the activation of NFκB pathway, which suggest that contribution from specific proteins to RT-EV-induced microglial activation could be occurring.

An additional explanation involves the presence of miRNAs that can modulate microglia. Previous research has shown that specific miRNAs have the ability to modulate oxidative stress, inflammatory responses, and consequently, cytokine release (Bala, Babuta et al. 2021). Moreover, some research confirmed that EVs carrying certain miRNAs can bind to TLR7 and activate microglia (Hu, Liao et al. 2018, Liao, Niu et al. 2020). Further experiments should aim to investigate the key molecule by which RT-EVs initially engage with the NFκB pathway.

Nevertheless, we showed that RT-EVs trigger the release of neurotoxic microglial H_2_O_2_. This is consistent with the well-studied neuronal death after H_2_O_2_ exposure (Hinshaw, Miller et al. 1993, Whittemore, Loo et al. 1995). It is important to note that the cell membrane is highly permeable to H_2_O_2_ and mediated transport also occurs, allowing for secreted H_2_O_2_ to go back into the cytosol and propagate redox signaling intracellularly. Additionally, the diffusion range of H_2_O_2_ is greater in the extracellular space. (Haslund-Vinding, McBean et al. 2017, Vilhardt, Haslund-Vinding et al. 2017). In combination, these autocrine and paracrine roles of H_2_O_2_ in microglia support not only the neurotoxic effect, but also the possibility of H_2_O_2_ propagating the initial oxidant effect both on the cell of origin and neighboring cells, altering their redox state and extending the inflammatory effect.

Our findings support a model where microglia drive neuroinflammatory responses. Nonetheless, other cells could contribute to the observed phenotype. Astrocytes are the most abundant cells in the brain, and they play a critical role in brain function (Yu, Wang et al. 2025). Even though microglia react faster and can influence astrocyte activation, studies focused on microglia-astrocyte cross talk found that astrocytes can activate microglia and both can modulate each other simultaneously and amplify neuroinflammatory responses (Yang, Wang et al. 2021, Candelario-Jalil, Dijkhuizen et al. 2022). Given this connection, it is plausible that astrocytes contribute to RT-EV-induced neuroinflammation, either directly or through interaction with factors released from microglia cells.

## 5. CONCLUSION

In summary, this study provides evidence that EVs derived from GBM after radiation treatment exert significant biological effects on microglial cells, demonstrating that RT-EVs can modulate brain function through oxidative and inflammatory responses. Mechanistically, we identify NFκB as a key pathway underlying RT-EV-mediated microglial activation, and knockdown of a critical subunit attenuated the neurotoxic effects of activated microglia. Our data show that NFκB is required for RT-EV-induced redox imbalance in microglia and may also contribute to the cognitive effects observed in vivo.

Together, our studies support a model where GBM-RT-EVs modulate brain homeostasis and provide insights into the role of EVs in redox modulation, cognition and suggest potential targets to treat GBM-associated cognitive impairment.

## 6. ACKNOWLEDGEMENTS

The authors would like to acknowledge the Animal Behavior Core (Robert Kline), Light Microscopy Core (Xu Fu), Pathology Research Core (Dana Napier), Biospecimen Procurement and Translational Pathology Shared Resource and Redox Metabolism Shared Resource Facility. Louis Rodgers, Malina Rijal and Cooper Kincaid for experimental assistance. This work was supported by: NCI: CCSG P30 CA177558, NCI: R01 CA251663 (LC.), NIH S10OD032256 for SARRP, NIGMS: CNS Metabolism P20 GM148326, NIGMS MCC-CCM P20 GM121327, 2023 Collaborative Bench to Bedside Radiation Medicine MCC, KY INBRE P20GM103436-24.

