## Supplementary for "Glioblastoma-derived extracellular vesicles released after radiation promote cognitive impairment through NFκB-mediated microglial activation"

**Methods**

Confirmation of delivery to the brain

To confirm delivery of fluorescently labeled EVs to the brain after intranasal administration, AMI HTX (Spectral Instruments Imaging) was used. AMI HTX is a system that allows for *in vivo* fluorescence and luminescence imaging. Mice were sedated with isoflurane, imaged and then sacrificed to harvest the brain which was imaged immediately after. Images were taken with an exposure of 15 s (excitation 573 nm, emission 588 nm)

**Figures**

**Supplementary Figure 1**

**A**

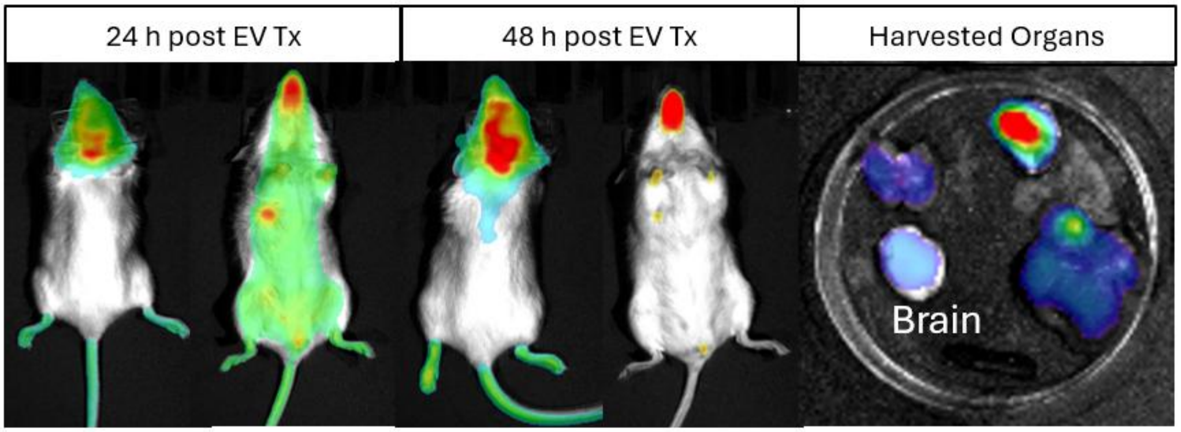
